# Perceptual decisions exhibit hallmarks of dynamic Bayesian inference

**DOI:** 10.1101/2022.05.23.493109

**Authors:** Julie A. Charlton, Wiktor F. Młynarski, Yoon H. Bai, Ann M. Hermundstad, Robbe L. T. Goris

## Abstract

To interpret the sensory environment, the brain combines ambiguous sensory measurements with context-specific prior experience. But environmental contexts can change abruptly and unpredictably, resulting in uncertainty about the current context. Here we address two questions: how should context-specific prior knowledge optimally guide the interpretation of sensory stimuli in changing environments, and do human decision-making strategies resemble this optimum? We probe these questions with a task in which subjects report the orientation of ambiguous visual stimuli that were drawn from three dynamically switching distributions, representing different environmental contexts. We derive predictions for an ideal Bayesian observer that leverages the statistical structure of the task to maximize decision accuracy and show that its decisions are biased by task context. The magnitude of this decision bias is not a fixed property of the sensory measurement but depends on the observer’s belief about the current context. The model therefore predicts that decision bias will grow with the reliability of the context cue, the stability of the environment, and with the number of trials since the last context switch. Analysis of human choice data validates all three predictions, providing evidence that the brain continuously updates probabilistic representations of the environment to best interpret an uncertain, ever-changing world.

**SIGNIFICANCE:** The brain relies on prior knowledge to make perceptual inferences when sensory information is ambiguous. However, when the environmental context changes, the appropriate prior knowledge often changes with it. Here, we develop a Bayesian observer model to investigate how to make optimal perceptual inferences when sensory information and environmental context are both uncertain. The behavioral signature of this strategy is a context-appropriate decision bias whose strength grows with the reliability of the context cue, the stability of the environment, and with the number of decisions since the most recent change in context. We identified exactly this pattern in the behavior of human subjects performing a dynamic orientation discrimination task. Together, our results suggest that the brain continuously updates probabilistic representations of the environment to make perceptual decisions in the face of uncertainty over both sensory and contextual information.

---

To accomplish goals, humans and other animals must infer properties of the environment in the face of uncertainty and change ^1,2^. Prior knowledge is often leveraged to guide perceptual decisions based upon ambiguous sensory measurements ^3^–^6^. However, knowledge that is relevant in one context may lead to worse outcomes if applied in another ^7^–^9^. A complex challenge arises when perceptual uncertainty is compounded by additional uncertainty about whether a change in context has occurred ^10,11^. As an example, imagine you are moving through a field, foraging for ripe bananas. Some bananas are clearly green or yellow and easy to judge, but many are ambiguous (Fig. 1a). Prior knowledge about the probability of encountering a ripe banana helps to make more accurate decisions. Bananas grown in sunny groves are more likely to be ripe, whereas those grown in shady groves are less likely to be ripe. As you move through the field with the sun overhead, it will be easy to identify the sunny and shady groves and use the appropriate prior knowledge. But clouds form, and the difference between sunny and shady groves becomes less clear. How can context-specific knowledge remain useful in the face of uncertainty over both perception and con-text?

**Figure 1.**
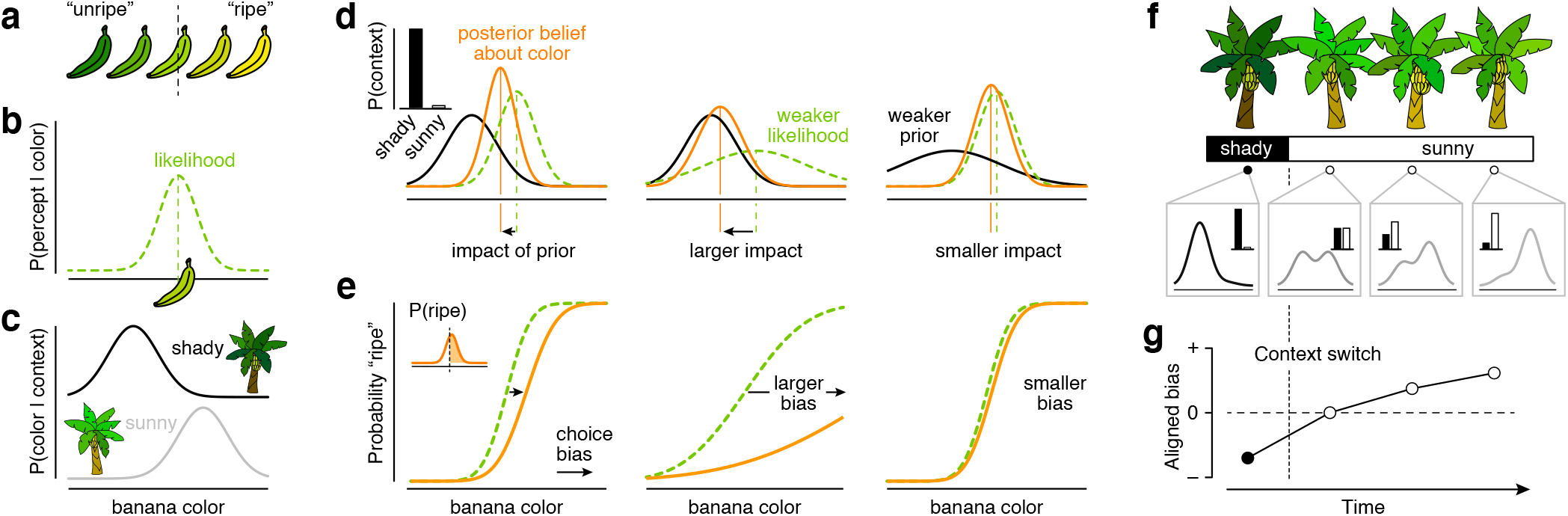
Example of Bayesian decision-making in a dynamic environment. (a) Yellower bananas are more likely to be ripe than greener bananas. (b) The likelihood distribution associated with a given color percept. (c) The conditional color distributions for bananas grown in shady vs sunny groves. (d) Illustration of a Bayesian foraging strategy. When the environmental context is certain (left, inset), the associated color distribution specifies the prior. The product of the prior and the likelihood gives the posterior. The impact of the prior on the posterior depends on the relative strength of likelihood and prior. (e) The forager decides that the banana is ripe when the probability that it is more yellow than some fixed criterion exceeds 50 percent (left, inset). The resulting choice behavior is plotted against the fruit’s color. The dotted green line illustrates this relation in the absence of prior knowledge, the full orange line illustrates this relation for the Bayesian forager. The prior biases the decision. (f) As the forager moves from one grove to another, she encounters changes in environmental context that are difficult to detect. When the context is uncertain, a Bayesian decision-maker constructs a prior by linearly mixing both color distributions, with weights determined by the continually-evolving context belief. (g) As a consequence, the influence of context-specific knowledge on choice behavior is minimal following a context transition, and grows over series of decisions uninterrupted by a context switch.

Bayesian inference offers a normative framework that specifies how knowledge can be optimally leveraged when making decisions under uncertainty ^12^. In the above example, knowledge about the ambiguity of perception is used to compute the likelihood of the perceived color given a banana’s true color (Fig. 1b), while knowledge about probable banana colors in shady and sunny groves is summarized as a context-specific prior belief (Fig. 1c). The normalized product of the prior and likelihood yields a posterior belief (Fig. 1d, left), which is used by a Bayesian decision-maker to decide whether or not to pick the banana (Fig. 1e, left). The impact of the prior on the decision depends on the relative strengths of the prior and the likelihood. When a sensory measurement is highly ambiguous (e.g., when assessing colors at dusk), the likelihood function is broad, and the same prior will have a larger impact on the posterior (Fig. 1d,e, middle panel). On the other hand, when the environmental context only weakly specifies the distribution of colors (e.g., in groves with mottled light), the prior is broad and will have a comparatively small impact (Fig. 1d,e, right panel).

Here, we develop a Bayesian ideal observer model to extend these normative predictions to a dynamic environment. Key to our predictions is the observer’s construction of a continually evolving posterior belief about the current context that informs the prior over stimuli (Fig. 1f). We find that this Bayesian strategy has three distinct signatures: First, when the context cue is more reliable, the observer is less uncertain about the identity of the current context, and will exhibit greater overall context-appropriate bias (hereafter, positive “aligned bias”). Second, when the environment is more stable (i.e., context switches happen less frequently), the observer will be overall less uncertain about the context, resulting in greater levels of aligned bias. Third, as an observer makes more decisions within the same context, they become less uncertain about the identity of the current context, and their aligned bias will grow (Fig. 1g).

We then asked whether human observers similarly leverage multiple forms of knowledge when making decisions in a dynamic environment. Subjects were shown a brief presentation of a drifting grating and asked to judge its orientation. Stimuli were drawn from one of three dynamically switching distributions, each representing a specific environmental context. At the beginning of each trial, an ambiguous cue (the color of the fixation mark) indicated the current context. Subjects were not told what this cue signified but had experienced the associated stimulus distributions in a prior training session. Analysis of the human choice behavior revealed a dynamically evolving influence of context that resembled the predictions of our Bayesian ideal observer. These results suggest that the brain continuously updates probabilistic hierarchical representations to combat the challenges posed by uncertain and unstable environments.

## RESULTS

Twelve human subjects performed a two-alternative forced choice (2AFC) orientation discrimination task in which they judged on every trial whether a visual stimulus presented in the near periphery was rotated clockwise or counterclockwise relative to vertical (Methods). Stimuli consisted of drifting gratings with variable orientation and contrast (Fig. 2a, top). Observers performed the task under three contexts characterized by different distributions of stimulus orientation (Fig. 2a, bottom). Context switches occurred pseudo-randomly. During the initial training phase, context switches were relatively rare, but they occurred frequently in the subsequent test phase (Methods). To quantify this aspect of the task, we computed for each trial the number of trials since the most recent context switch. This metric was approximately exponentially distributed across trials and had a mean value of 15.9 trials during the training phase and 2.35 trials during the test phase (Fig. 2b).

**Figure 2.**
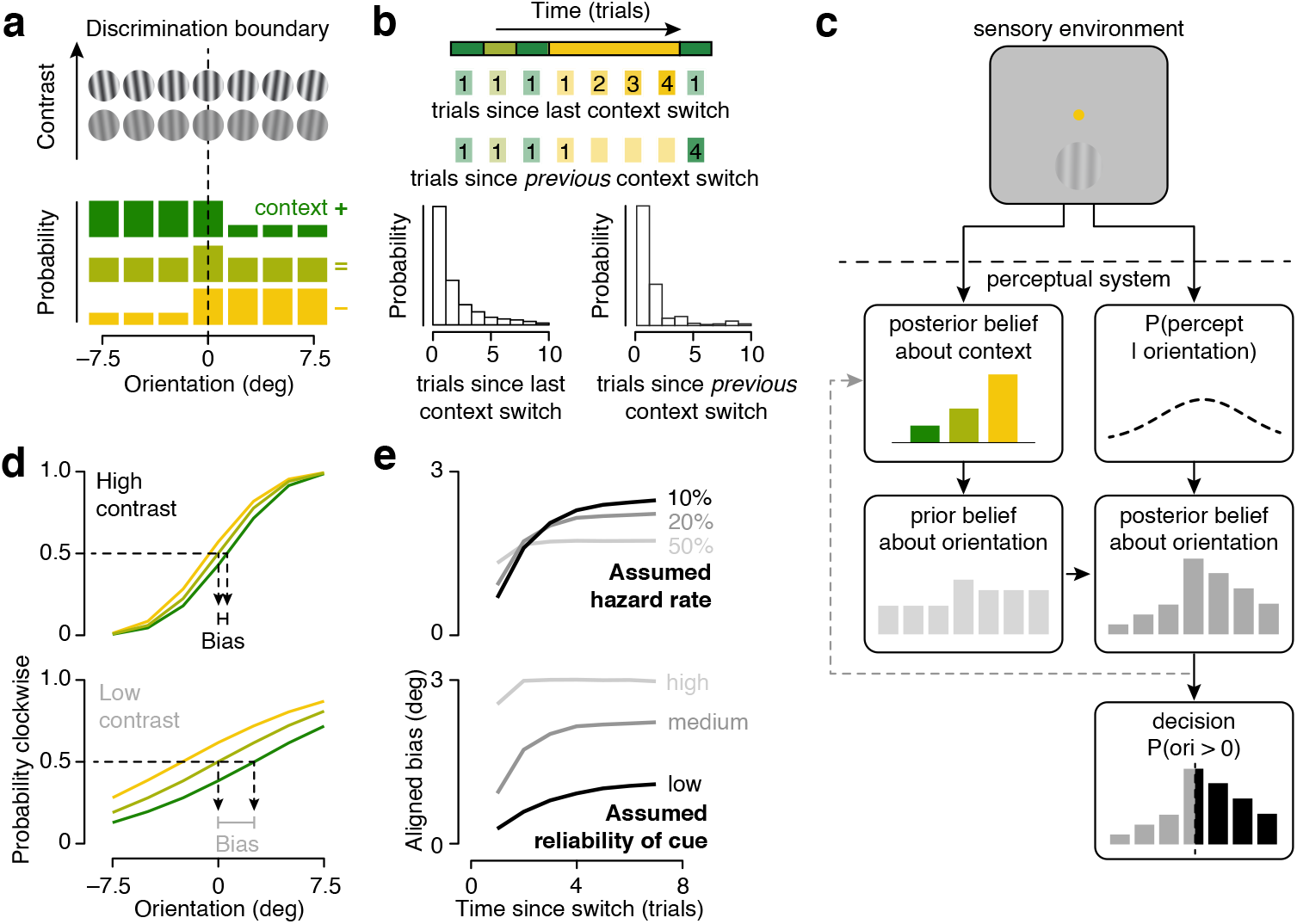
Bayesian ideal observer model for dynamic 2AFC orientation discrimination. (a) Context-specific distribution of stimulus orientation used in the model simulations. Stimulus contrast was varied over two levels. (b) The cross-trial distribution of the number of trials from the most recent context switch. (c) Schematic of the Bayesian ideal observer’s decision-making process. (d) Summary of the model’s choice behavior. Proportion of clockwise choices is plotted against stimulus orientation, split out by task context (different lines) and stimulus contrast (top vs bottom). (e) Evolution of low contrast aligned bias as a function of the number of trials since a context switch split out by assumed hazard rate (top) and assumed reliability of the context cue (bottom).

This task is difficult for several different reasons. First, stimulus strength (i.e., rotation magnitude) is weak in light of perceptual acuity for orientation ^13^,14. Second, at low contrast, perceptual sensitivity is further reduced, and uncertainty about stimulus orientation is elevated ^15,16^. And third, while there are context-specific regularities that can be leveraged to improve performance (i.e., two out of three contexts are associated with a skewed distribution of orientations), the environment is constantly in flux, leaving observers potentially uncertain about the underlying context on any given trial. As outlined above, a Bayesian decision-maker faced with these challenges maximizes performance by exploiting its knowledge of the statistical structure of the task. In the following section, we use this general concept to formulate a set of specific predictions for our task.

### Bayesian ideal observer model for dynamic orientation discrimination

To identify signatures of the optimal decision-making strategy for this task, we investigated the choice behavior of a Bayesian ideal observer model. Like our human subjects, the ideal observer was presented with a sequence of oriented stimuli and tasked to decide on every trial whether a given stimulus was rotated clockwise or counter-clockwise. It does so on the basis of both a noisy measurement of the stimulus orientation and an ambiguous context cue (Fig. 2c, top). Specifically, the ideal observer computes the likelihood of the sensory measurement given the stimulus’ true orientation and combines this with its prior belief about stimulus orientation to obtain a posterior belief about orientation (Fig. 2c, middle), which in turn informs the decision (Fig. 2c, bottom).

The ideal observer assumes that the environment switches between three discrete contexts at a fixed rate, and it knows the underlying distribution of stimuli in each context. Thus, if there were no uncertainty about the current context, the ideal observer would use the appropriate context-specific stimulus distribution to help disambiguate incoming stimuli. However, the ideal observer also assumes that the context cue itself is ambiguous. As a result, it must use the incoming context cue, together with previously decoded stimuli, to continually update its posterior belief about context (Methods, Fig. 2c, middle). The ideal observer then uses this posterior belief over contexts to update its prior belief about stimulus orientation for the current trial.

The inference strategy manifests in patterns of decisions that cannot be deduced from any individual choice, but rather become evident in the relationship between choice and task variables. To expose this relationship, we simulated a large number of trials in our task. Plotting average choice behavior as a function of stimulus orientation, split out by underlying task-context, reveals that the ideal observer’s decisions depend both on sensory stimulus measurements and context-specific prior knowledge (Fig. 2d). The influence of context on the decision depends on the relative strength of the prior and likelihood. For example, in low contrast settings, the sensory orientation measurement is more uncertain (i.e., the likelihood is broader), and the association between stimulus orientation and behavioral choice is weaker, as evidenced by the shallower slope of the lines. The ideal observer naturally compensates for this loss of information by relying more heavily on context-specific knowledge. This results in a stronger aligned bias, as evidenced by the increased horizontal separation between the lines (Fig. 2d). We quantify the influence of context on the decision by computing the average change in the point of subjective equality (defined as the orientation that elicits 50% clockwise choices) from the uniform context.

When the context changes over time, the prior over stimuli will naturally weaken and strengthen as a function of the ideal observer’s evolving belief about context. Following a context switch, incoming stimuli and context cues are in conflict with the ideal observer’s belief about context, which leads to an increase in uncertainty about context and a weakening of the context posterior. This, in turn, leads to a weakening of the stimulus prior, which will evolve from a single context-specific distribution to a mixture distribution (see example in Fig. 2c, middle). In this way, our Bayesian ideal observer will at times use a prior that *does not* correspond to any individual context. Weakening of the prior due to uncertainty about context is evident in the evolution of the ideal observer’s aligned bias as it completes more trials within a context. Fig. 2e shows the temporal evolution of aligned bias in low contrast settings for different assumptions adopted by the ideal observer about the exact statistical structure of the task. Consider the overall trend. Just after a context switch (i.e., a low stability level), the ideal observer has high uncertainty about the current context, and the context-induced bias is minimal. As the ideal observer performs more trials within a context, it continually updates its belief and reduces its uncertainty about the current context, and the aligned bias grows.

The particular pattern of aligned bias depends on the ideal observer’s underlying assumptions about the stability of the environment and the reliability of the context cue. Aligned bias evolves dynamically whenever the ideal observer assumes the environment to be somewhat stable (i.e., the assumed probability of a context switch is less than 0.5; Fig. 2e, top) and the context cue to be ambiguous (Fig. 2e, bottom). The higher the assumed stability of the environment and the lower the assumed reliability of the context cue, the longer it takes for the aligned bias to fully saturate (Fig. 2e). The level at which the aligned bias saturates is higher when context changes are assumed to be less probable and the cue more reliable (Fig. 2e). The temporal evolution of the aligned bias also depends on the number of trials since the *previous* context switch. The more stable the environment is, the stronger the ideal observer’s belief is about the previous context, and thus the more evidence (time) that is required to update that belief and the longer it takes for the context to maximally exert its influence on choice (Fig. SS1a). However, as can be appreciated from the subtle horizontal shift in color across the rows of the bias-matrices in Fig. SS1b, this effect is generally weak compared to the overall influence of context stability. Finally, note that even in the most extreme cases, the aligned bias does not go negative. In principle, this could happen. However, under the specific conditions we studied, it does not.

In summary, a Bayesian ideal observer uses a hierarchical inference strategy to perform our task. This yields orientation judgments that are biased by task context. The magnitude of this effect not only depends on stimulus contrast, but also on the observer’s belief about the reliability of the context cue, the stability of the environment, and on the number of trials since the most recent context switch.

### Effect of cue reliability, context volatility, and sensory uncertainty on human choice behavior

What knowledge do humans leverage when making perceptual decisions in dynamic environments? Leveraging knowledge about context-specific priors and perceptual ambiguity biases uncertain perceptual decisions ^3^–^9^. As we have shown, additionally leveraging knowledge about the hierarchical structure of our task and the stability of the environment should minimize the magnitude of this bias right after a context switch. Moreover, this effect should also depend on the reliability of the context cue and sensory uncertainty. To test these predictions, we manipulated critical task statistics and conducted several targeted analyses of the human choice behavior. To study the effect of the reliability of the context cue, we assigned each subject to one of two conditions. In the veridical cue condition, the context cue correctly indicated the underlying context on every single trial. In the ambiguous cue condition, the cue was valid on 80% of the trials. Subjects were not told what the cue signified, but experienced the associated stimulus distributions during the initial training phase (Fig. 3a). To study the effect of the stability of the environment, context switches were relatively rare during the training phase, but occurred frequently in the subsequent test phase (Fig. 4a, top).

**Figure 3.**
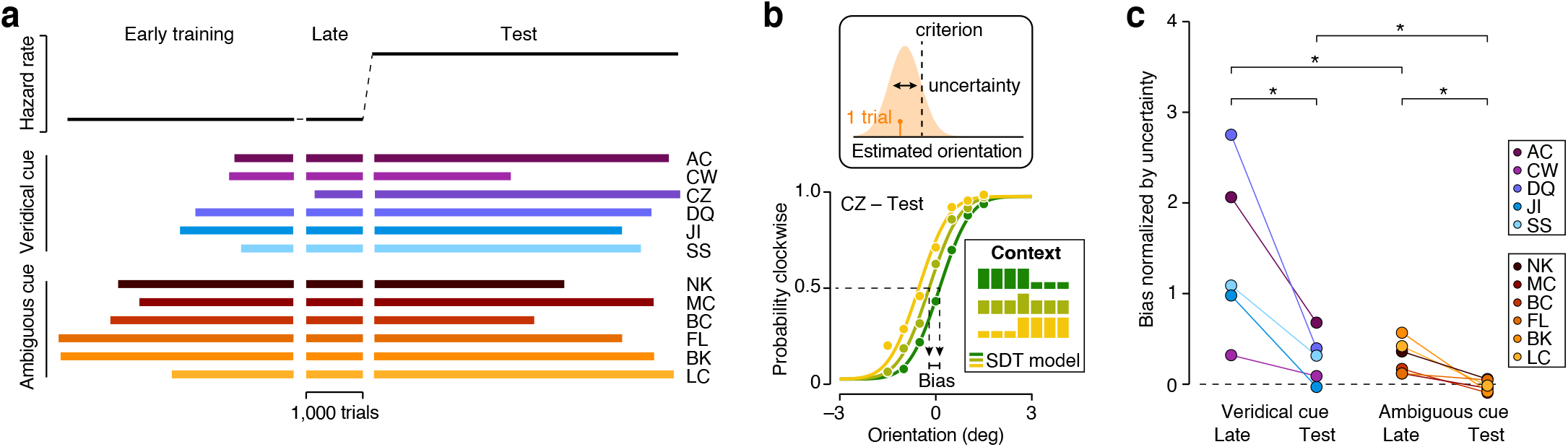
Human orientation judgments are biased by task-context in a manner that depends on the reliability of the context cue and on the stability of the environment. (a) Number of completed trials during the training and test phase of the experiment for each subject. (b) Bottom: Proportion of clockwise choices of an example observer plotted as a function of stimulus orientation under three dynamically-switching task contexts (indicated by color). Symbols summarize observed choice behavior, lines show the fit of a Signal Detection Theory model of decision-making. Top: In the model, a stimulus gives rise to a noisy orientation estimate. Comparison with a fixed criterion yields a decision. (c) Comparison of normalized bias during the training and test phase. In the veridical cue condition, cue reliability was 100%; in the ambiguous cue condition, cue reliability was 80%. * P < 0.05.

We first asked whether choices were biased by task context in a manner that depends on the reliability of the context cue and the stability of the environment. Consider an example human observer who judges the same stimuli differently under different contexts during the test phase (Fig. 3b, bottom panel). To quantify this effect, we described the data with a Signal Detection Theory (SDT) based process-model of decision-making that specifies how the probability of a “clockwise” choice depends on the task variables (orientation, contrast, and context; lines in Fig. 3b, bottom panel). We then used this model to measure the observers’ uncertainty about stimulus orientation (defined as the cross-trial variability in the orientation estimate, Fig. 3b, top panel) and the magnitude of their aligned bias. Dividing this latter statistic by the former provides a normalized estimate of aligned bias. For each subject, we independently estimated their aligned bias at the end of the training phase and during the test phase. We only included trials of the same contrast level in this analysis (see Methods). As can be seen in Figure 3c, the sudden decrease in the stability of the environment led to decreased aligned bias for every single subject (median decrease in normalized bias = 0.435, P < 0.001, one-sided Wilcoxon signed-rank test, n = 11). This effect was significant within each condition (veridical cue condition: median decrease = 1.01, P = 0.031, n = 5; ambiguous cue condition: median decrease = 0.283, P = 0.016, n = 6). Recall that the context cue was less reliable in the ambiguous cue condition than in the veridical cue condition. As predicted, this resulted in lower levels of aligned bias (Fig. 3c). This was true both at the end of the training phase (median bias = 1.084 for the veridical cue condition and 0.259 for the ambiguous cue condition, P = 0.015, one-sided Wilcoxon rank-sum test), and during the test phase (median bias = 0.31 for the veridical cue condition and –0.035 for the ambiguous cue condition, P = 0.015). We conclude that aligned bias increases as the context cue becomes more reliable, but decreases as the environment becomes less stable.

A hallmark of Bayesian inference is the stronger reliance on context-specific knowledge when sensory measurements are more uncertain. To test whether human choices exhibit a similar pattern, we next asked whether choices were biased by task context in a contrast-dependent manner. During the test phase, high and low contrast stimuli were pseudo-randomly intermixed (see Methods). For each subject, we independently estimated their uncertainty about stimulus orientation and the magnitude of their aligned bias for high and low contrast stimuli. As can be seen in Figure 4a, lowering stimulus contrast increased orientation uncertainty for all subjects but one (median increase in uncertainty = 0.303 deg, P < 0.001, one-sided Wilcoxon signed-rank test, n = 12). This effect was significant within each condition (veridical cue condition: median increase = 0.403 deg, P = 0.016, n = 6; ambiguous cue condition: median increase = 0.147 deg, P = 0.031, n = 6). In the veridical cue condition, lowering stimulus contrast also resulted in a larger aligned bias, though note that one subject (JI) did not exhibit context-dependent choice behavior at either high or low contrast (median increase = 0.248 deg, P = 0.031, n = 6; Fig. 4b, left panel). In the ambiguous cue condition, there was no consistent aligned bias during the test phase of the experiment (high contrast: median bias = –0.032 deg, P = 0.989, n = 6; low contrast: median bias = –0.019 deg, P = 0.784, n = 6), nor a consistent change in the magnitude of the bias (median increase = 0.018 deg, P = 0.156, n = 6; Fig. 4b, right panel). Thus, when present, aligned bias increases with stimulus uncertainty.

**Figure 4.**
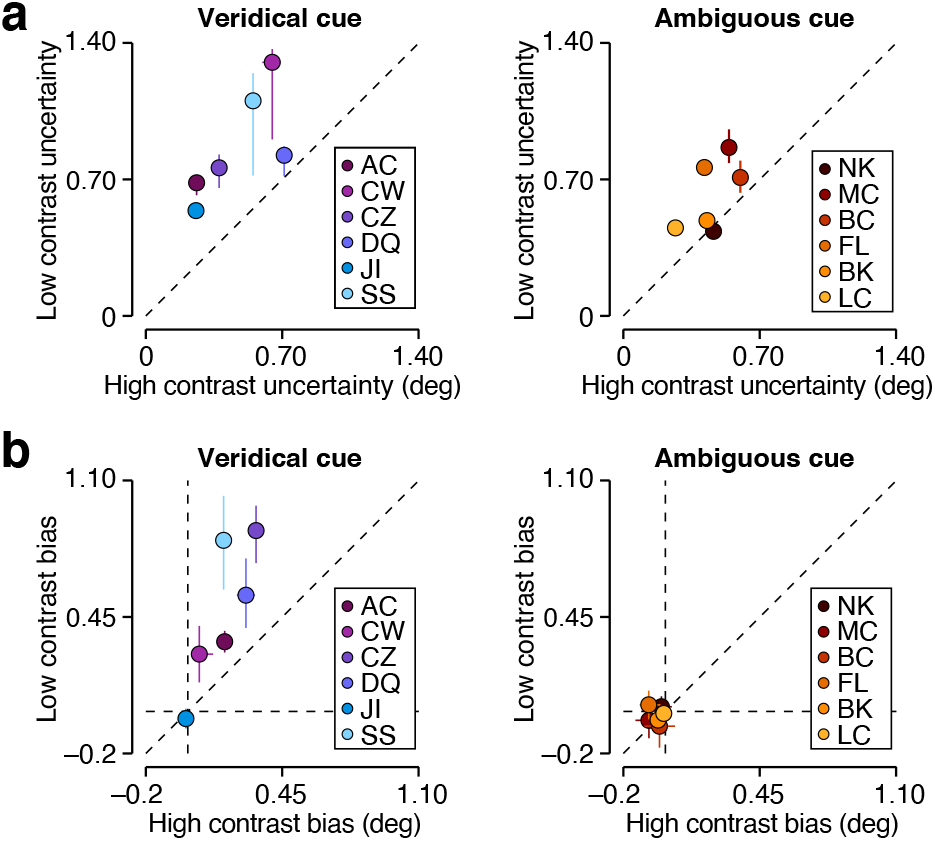
Human orientation judgments are biased by task-context in a contrast-dependent manner. (a) Orientation uncertainty for low (ordinate) and high (abscissa) stimulus contrasts for the veridical cue condition (left) and the ambiguous cue condition (right) subjects. (b) Aligned bias for low (ordinate) and high (abscissa) stimulus contrasts for the veridical cue condition (left) and the ambiguous cue condition (right) subjects. Error bars reflect the 68 percent confidence interval, derived from a 1,000-fold bootstrap analysis (Methods).

So far, we have shown that orientation judgments are biased by task context. The magnitude of this effect depends on stimulus contrast and the reliability of the context cue. This suggests that subjects in our task typically incorporate knowledge about perceptual ambiguity and context-specific priors into their decision-making process. The magnitude of the aligned bias also depends on the stability of the environment. This suggests that observers additionally exploit knowledge about the hierarchical structure of the task and the stability of the environment. All these effects support the hypothesis that subjects continuously update a probabilistic model of the environment to interpret ambiguous sensory stimuli. To further test this hypothesis, we now turn to the question of whether aligned bias changes with the number of trials since the most recent context switch (i.e., time spent in the current context).

### Effect of trials since a context switch on human choice behavior

Does the influence of context on the decision change over time following a context switch? This is a difficult question to address for two reasons. First, we have a limited amount of choice data (mean = 4,120 completed trials during the test phase per observer), distributed unevenly across waiting times (Fig. 2b). Estimating aligned bias separately for each “level” of trial count post-context switch, as we did for the ideal observer, would yield unreliable estimates for our human observers. In other words, the frequency of context switches in our experiment creates an abundance of trials that occur right after a context switch and many fewer trials that are, for example, the tenth trial within a single context. Second, in perceptual decision-making tasks, observers commonly exhibit sequential choice dependencies that here could be mistaken for Bayesian-like dynamic inference. Specifically, observers’ responses are often correlated with their previous response ^17^ and, sometimes, with the previous stimulus ^18^. These correlations are usually positive, but can be negative as well. Although such dependencies may improve overall decision accuracy in temporally continuous environments, they are distinct from dynamic Bayesian inference, which relies on the continual updating of one’s belief about context.

To overcome these challenges, we developed a descriptive modeling approach to characterize the evolution of context-specific aligned biases following a context switch. Our methodology is related to approaches developed by Roy et al. (2021) ^19^ to characterize the temporal evolution of decision-making strategies in static environments. Specifically, we use a dynamic Bernoulli generalized linear model (GLM) defined by a set of weights that specify the trial-by-trial influence of different task variables on the observer’s decision. These variables capture stimulus and context manipulations, as well as recent response and stimulus history (Fig. 5a). Our approach is novel in its use of a dynamic “bias function”, *f* (*S*), that describes the temporally-evolving influence of context on choice and is given by:

**Figure 5.**
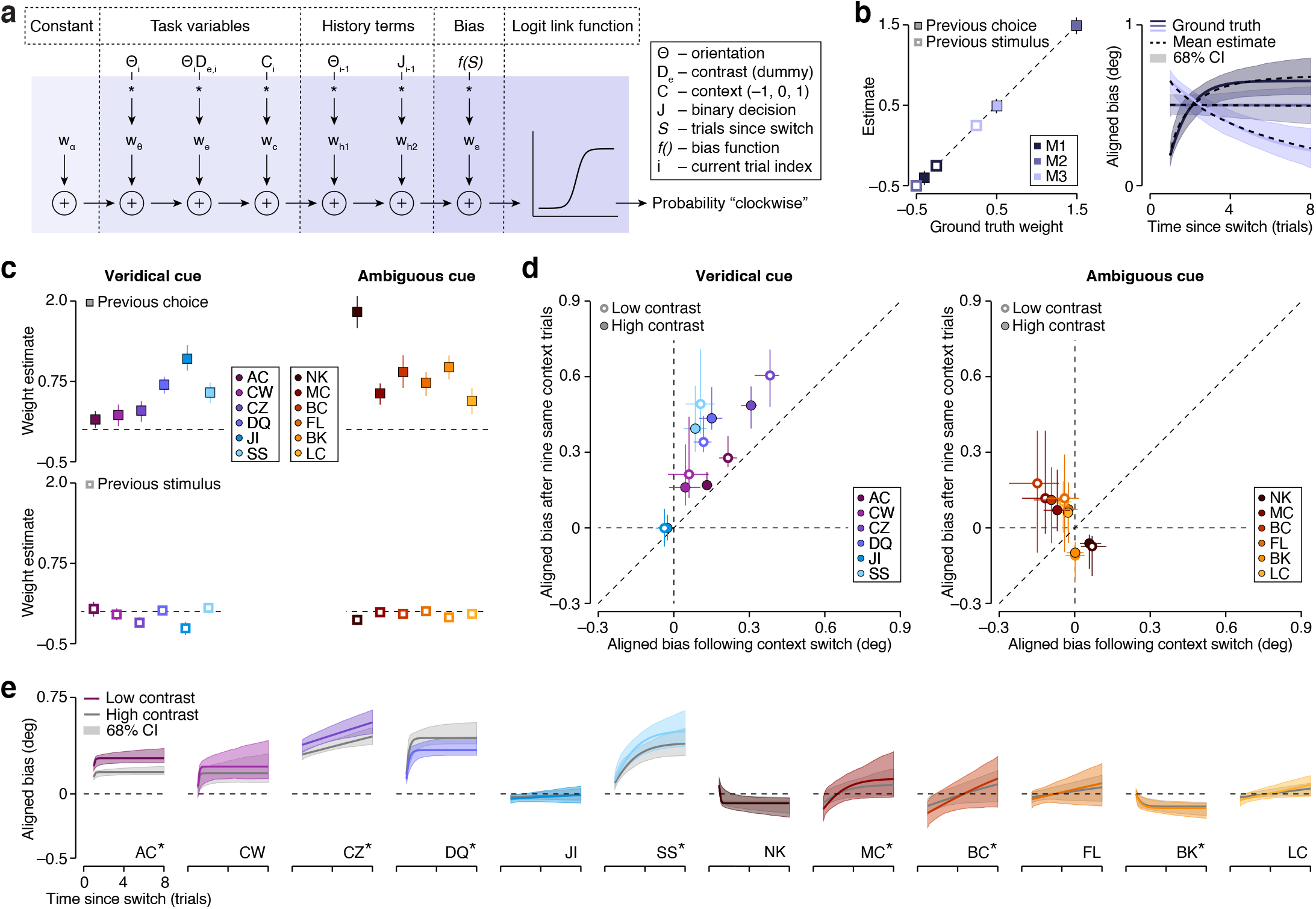
GLM-based analysis of cross-trial dynamics in human orientation judgment strategies. (a) Dynamic GLM. The probability of a clockwise choice is predicted by the logistic transformation of six linearly combined regressors. (b) Recovery analysis. Left: Estimated history terms plotted as a function of their ground truth value for three model observers. Symbols indicate the mean value; error bars, where visible, illustrate the 68 percent confidence interval. Right: Evolution of aligned bias for high contrast stimuli as a function of the number of trials since the most recent context switch for three model observers. The colored line indicates the ground truth relationship, the dotted black line shows the mean estimate, and the shaded region illustrates the 68 percent confidence interval. (c) Estimated history terms for veridical cue condition (left) and ambiguous cue condition (right) subjects. (d) Model-predicted bias after ten same-context trials plotted against bias immediately following a context switch for the veridical cue condition (left) and the ambiguous cue condition (right) subjects. Error bars illustrate the 68 percent confidence interval. (e) Evolution of aligned bias for low and high contrast stimuli as a function of trials since a context switch for all subjects. The shaded region illustrates the 68 percent confidence interval. *Dynamic GLM outperforms static GLM in cross-validation analysis (see Supplementary Table S4). All confidence intervals were computed from 1,000 simulated data sets.

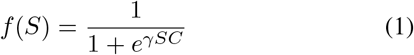

where *S* is the number of trials since the last context switch (for the current trial), *γ* controls the shape of the bias function, and *C* is a categorical variable that takes a value of –1, 0, or 1 for the negatively skewed, uniform, or positively skewed context. We chose this functional form because it can capture a variety of monotonically evolving relationships. For each observer, we jointly estimate the weights and the shape parameter *γ* by maximizing the likelihood of the data under the model. Because *f* (*S*) varies nonlinearly with *S*, we use a two-step grid-search procedure to find this maximum (Methods).

To validate our method, we used the dynamic GLM to generate synthetic data sets for three model observers. These model observers had identical weights for the task variables, but differed in their reliance on past history and in the evolution of their aligned bias with the number of trials following a context switch. Each model observer was presented with the exact sequence of trials presented to one of our human observers. We then applied our analysis procedure to these synthetic data, and we found that it provided a robust and unbiased estimate of the influence of response and stimulus history (Fig. 5b, left panel), as well as of the dynamic influence of the number of trials since the last context switch. This latter point can be best appreciated by considering recovery of the evolution of the bias function (Fig. 5b, right panel). We obtain this relation by setting the model’s history terms to zero, such that:

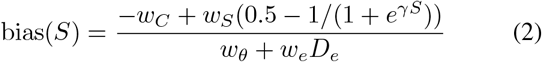

where *w*_*C*_ is the weight on context, *w*_*S*_ the weight on the bias function, *w*_*θ*_ the weight specifying the effect of stimulus orientation at low contrast, *w*_*e*_ the weight specifying the additional effect of orientation at high contrast, and *D*_*e*_ a dummy variable that takes a value of 0 for low contrast stimuli and 1 for high contrast stimuli. As can be seen in Fig. 5b, the fitted GLM closely approximates the ground truth effects of dynamic bias. This is true for a simulated aligned bias that rapidly rises, is independent of, or slowly decreases with the number of trials since the last context switch.

Having validated our method, we described each human observer’s choice behavior with the dynamic GLM and used a bootstrap-based procedure to obtain confidence intervals for the model predictions (Methods). The dynamic GLM has only one more free parameter than the static SDT-model, but described the data much better (Fig. S2, see Supplementary Table S1 for AIC comparison). This improvement was in part due to the inclusion of history terms. In particular, we found that observers’ responses were systematically correlated with their previous response (Fig. 5c, top), but not with the previous stimulus (Fig. 5c, bottom). In addition, the use of a dynamic bias function helped to capture the changing influence of context on the perceptual decision following a context switch. A cross-validation analysis revealed that this model component was necessary for some, but not all, subjects (see Supplementary Table S4). In spite of this heterogeneity, we robustly observe that the influence of context on the perceptual decision grows with time spent in the current context. To quantify this effect, we calculated the model-predicted bias for trials that occurred ten trials after a context switch (Eq. 2) and compared this value to the model’s prediction for trials that immediately followed a context switch. As can be seen in Figure 5d, spending ten consecutive trials in the same context increased aligned bias for most subjects (median increase in aligned bias at high contrast = 0.107 deg, P = 0.005, one-sided Wilcoxon signed-rank test, n = 12; low contrast = 0.155 deg, P = 0.017, n = 12). This effect reached statistical significance within the veridical cue condition (high contrast = 0.147 deg, P = 0.016, n = 6; low contrast = 0.188 deg, P = 0.016, n = 6), but not within the ambiguous cue condition (high contrast = 0.094 deg, P = 0.281, n = 6; low contrast = 0.154 deg, P = 0.078, n = 6), perhaps due to the substantially weaker aligned bias within this condition. The estimated cross-trial evolution of the aligned bias is plotted for each subject in Figure 5e. Inspection of these curves reveals that for some subjects, aligned bias increased gradually over the course of multiple trials (CZ, SS, BC) while for others, the bias abruptly saturated (AC, CW, DQ), or barely changed at all (JI, FL, LC). Within the veridical cue condition, this inter-observer variability could be explained by our Bayesian ideal observer with a miscalibrated assumption of the stability of the environment or the reliability of the context cue. Note that in the ambiguous cue condition, several observers exhibit negative aligned bias either immediately after a context switch, or after multiple trials within a context. While this effect is not predicted by the Bayesian ideal observer, we suspect it was induced by the low environmental stability of the test phase as all ambiguous cue condition observers had positive aligned bias during the highly stable training phase (Fig. 3c). These results therefore suggest that our subjects’ decisions were not only guided by knowledge about perceptual ambiguity and context-specific priors, but also by knowledge about the hierarchical nature of the task and the stability of the environment.

## DISCUSSION

Perception is inherently uncertain, and natural environments are constantly in flux. Both factors are considered major forces that shape computation and representation in sensory systems ^16,20^–^24^. This raises the question of how uncertainty and instability jointly impact sensory-guided decisions ^11,25^. Our analysis reveals that in a simple perceptual decision-making task, human subjects rely more strongly on context-specific knowledge when their belief about the current context is more certain, which occurs in more stable environments and after spending more time in the same context. This strategy can be understood as a rational adaptation to the challenges presented by a dynamic world. Moments of transition often yield uncertainty about the underlying context. Does the first clap of thunder really announce the arrival of rain? Does the first flower truly signify that Spring has begun? The answer to these questions will impact subsequent decisions, but because the available cues are ambiguous, it is not possible to answer them correctly all of the time. The most accurate strategy is to build probabilistic hierarchical representations to guide the interpretation of incoming stimuli. Note that we do not explicitly model how observers learn these representations, though understanding which computations humans use to infer statistical properties of dynamic environments is an active area of research ^26–28^. Under this strategy, uncertainty about the current context weakens strong context-specific priors over stimuli. When newly incoming evidence (the sound of rain drops or the sight of melting snow) further clarifies the context, these priors –and their impact on perceptual decisions– grow stronger again. Our study complements recent work on uncertain perceptual decisions in dynamic environments. One set of studies asked how negative feedback influences decision-making strategies when stimulus-response contingency rules undergo covert and unpredictable changes, and found that both humans and monkeys take expected choice accuracy into account to disambiguate the source of failure ^29,30^. Another set of studies investigated temporal integration of dynamic stimuli in environments that change on different timescales. Human observers adapt their decision-making strategies to fluctuations that occur suddenly within a single trial ^31^, between trials in a sequence ^32^, or gradually across long blocks of trials ^33^. They do so in various manners, which include adjusting the rate at which they discount previous beliefs ^31^, biasing initial beliefs ^29^, and adopting timevarying decision criteria ^34^. To a first approximation, these various modifications can all be understood as attempts to implement the optimal Bayesian decision-making policy ^25^.

We found that uncertain perceptual decisions in changing environments are biased by context-specific knowledge in a manner that qualitatively resembles the ideal Bayesian strategy, but quantitatively deviates from this optimum. Specifically, we reported that decision bias is larger when context cues are more reliable (Fig. 3c, veridical vs. ambiguous cue condition), when the environment is more stable (Fig. 3c, late vs test), when the sensory measurement is less certain (Fig. 4b), and after more time has been spent in the current context (Fig. 5d). These effects are all predicted under a hierarchical Bayesian inference strategy (Fig. 2d-e). However, our analysis also reveals several quantitative deviations from the optimal strategy. Most notably, when the context cue is 100% reliable, there is no uncertainty about the current context under a well-calibrated generative model of the task, and hence decision bias should depend on neither the stability of the environment nor the number of trials since the most recent context switch. This is not what we observed in the veridical cue condition. Moreover, when the context cue is 80% reliable, a well-calibrated model would result in *some* decision bias, even in highly unstable environments. This is not what we observed during the test phase of the ambiguous cue condition. To a first approximation, subjects in both conditions behaved as if they used a Bayesian inference strategy, but relied on a mis-calibrated generative model of the task in which the reliability of the context cue was systematically underestimated.

The perceptual inference problems faced by humans and other animals are complex, and so are the strategies they use to make behavioral choices. Normative models are a critical tool to uncover the principles that shape these strategies. In our task, the ideal Bayesian strategy yields a dynamically evolving prior over stimuli. This prediction is difficult to test: the normative model is too complex to directly fit to choice data, but a pure data-based trial-averaging approach is not efficient enough to reliably characterize the temporal evolution of aligned bias from realistic amounts of data. Instead, we opted to test the normative prediction by looking at our data through the lens of different modeling approaches. We used a tried-and-tested process model of decision-making to verify the presence of uncertainty-dependent aligned biases and a flexible descriptive model to characterize the dynamic evolution of these biases. The interpretation of our data critically relies on the combined insights offered by each of these approaches. As such, our study provides an example of how different computational tools can be used in conjunction to gain insight into the mechanisms underlying complex choice behavior at the single trial level.

## METHODS

### Behavioral task

Twelve human subjects (5 male, 7 female; ages 18-30) with normal or corrected-to-normal vision participated in the experiment. The experimental protocol was approved by the local ethics committee (Institutional Review Board of The University of Texas at Austin) and all participants gave informed consent. Subjects were not aware of the purpose of the study. Owing to sample size, no gender-specific analyses were performed.

Subjects were seated in a dimly lit room in front of a gamma-corrected CRT monitor (Hewlett Packard, A7217A). A head and chin rest ensured that the distance between the participants’ eyes and the monitor’s screen was 57 cm. Eye position was recorded with a high-speed, high-precision eye tracking system (EyeLink 1000). We presented visual stimuli at a spatial resolution of 1280 × 1024 pixels and a refresh rate of 75 Hz. Stimuli were presented using PLDAPS software (https://github.com/huklab/PLDAPS) on an Apple Macintosh computer.

Subjects performed an orientation discrimination task in blocks of 48 trials. Each trial began when participants fixated a small square (0.5^°^diameter) at the center of the screen. After 500 ms, two choice targets appeared, one on each side of the fixation point (on the horizontal meridian, at 4.5 degrees eccentricity). The choice targets were white lines (2^°^long, 0.3^°^wide), rotated –22.5^°^(choice target on the left) and 22.5^°^(choice target on the right) from vertical. After 500 ms, a circularly vignetted drifting grating appeared. The stimulus was positioned in the lower left visual quadrant (centered at an eccentricity of 3.2^°^), measured 1.25^°^in diameter, had a spatial frequency of 2.5 cycles/deg, and a temporal frequency of 3 cycles/s. Subjects judged the orientation of the stimulus relative to vertical. The stimulus remained on for 500 ms. The stimulus then disappeared along with the fixation mark and subjects reported their decision with a saccadic eye movement to the choice target whose orientation was closest to the estimated stimulus orientation. Auditory feedback about the accuracy of the response was given at the end of each trial. We varied stimulus orientation over a small range (a few degrees) that was centered on vertical and tailored to each observer’s orientation sensitivity. Vertically oriented stimuli received random feedback. Stimuli were presented at either high or low contrast (Michelson contrast of 100% and 10%). For nine out of twelve observers, high and low contrast stimuli were randomly interleaved. For the three remaining observers, high and low contrast stimuli were grouped in blocks of eight trials.

Subjects performed the task under three contexts, characterized by a uniform, negatively skewed, and positively skewed distribution of stimulus orientation (shown in Fig. 2a). The corresponding baseline probability of a “clockwise” choice being correct was 50%, 70%, and 30%. Context switches occurred pseudo-randomly, with a hazard rate of 6.3% during the training phase, and of 42.5% during the test phase. The color of the fixation mark (red, green, or blue) also varied across trials. In the veridical cue condition, this color indicated the underlying context on 100% of trials, in the ambiguous cue condition, this was the case on 80% of the trials. Subjects were not told what the color of the fixation mark signified. Trials in which the subject did not maintain fixation within 1.5^°^of the fixation mark were aborted. Participants performed the task across three to eight sessions and successfully completed between 2,420 and 5,388 trials during the test phase of the experiment.

Before performing the main task, subjects participated in one or more training sessions. Compared to the main task, the range of orientations was larger and context switches occurred much less frequently in these training sessions. We considered subjects ready for the main task once their stimulus judgements were consistent (stable, lawfully shaped psychometric functions), reliable (few lapses on easy catch trials), and appropriately biased by context in the most difficult conditions. This typically required more than 2,000 training trials (see Fig. 3a).

### Bayesian ideal observer model

We derived predictions for a Bayesian ideal observer who leverages its knowledge of the statistical structure of the task to maximize decision accuracy. The ideal observer assumes that on each trial *t*, the stimulus orientation *θ*_*t*_ and the context cue *x*_*t*_ depend on the true underlying context *C*_*t*_, and that each context is associated with a specific stimulus distribution *p*(*θ*_*t*_ | *C*_*t*_) that is matched to one of the distributions used in the behavioral task (Fig. 2a). It further assumes that context switches occur with a probability *h* (i.e., the hazard rate), inducing the following transition probability over the context variable *C*_*t*_:

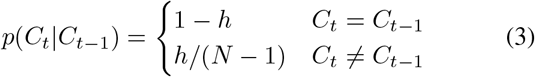

where *N* = 3 is the number of possible contexts.

On each trial, the ideal observer obtains a noisy orientation measurement *y*_*t*_ via the encoding distribution *p*(*y*_*t*_ | *θ*_*t*_, *σ*^2^) = 𝒩 (*θ*_*t*_, *σ*^2^), where *σ*^2^ is the variance of sensory noise. It also obtains a noiseless context cue measurement *x*_*t*_. The ideal observer then uses these measurements to update its beliefs about task context and stimulus orientation and make a decision.

We consider two ways in which the ideal observer’s generative model of the task can be miscalibrated: either the assumed hazard rate or the assumed reliability of the context cue (or both) may differ from the actual values used in the behavioral experiment (Fig. 2e). In particular, the ideal observer assumes that the context cue is perturbed by external noise via the cue-generating distribution 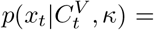, where 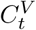 is the angular label of the context *C*_*t*_, and *κ* is the concentration parameter that controls the assumed reliability of the context cue. This formulation captures the situation whereby the first context could be mistaken for the third one and vice-versa.

To make a decision, the ideal observer estimates the probability *p*(*θ*_*t*_ *>* 0 | *y*_*t*_, *x*_*t*_) that the stimulus is oriented clockwise. This estimate is based on the following sequence of steps (note that for notational simplicity, we denote only the most recent stimulus *θ*_*t*_ and cue *x*_*t*_, instead of their respective histories *θ*_*τ*≤*t*_ and *x*_*t*≤*t*_):

1. Update the posterior over contexts with the measured context cue; i.e., compute *p*(*C*_*t*_|*x*_*t*_) *∝ p*(*x*_*t*_|*C*_*t*_)*p*(*C*_*t*_|*x*_*t*−1_)
2. Compute the prior over stimuli by marginalising over the posterior over contexts; i.e., compute 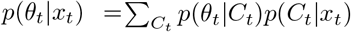
3. Use the resulting prior to compute the posterior over stimulus orientations *θ*_*t*_ given the noisy representation *y*_*t*_; i.e., compute *p*(*θ*_*t*_|*y*_*t*_, *x*_*t*_)
4. Use the resulting posterior to compute the probability that the stimulus is oriented clockwise; i.e., compute 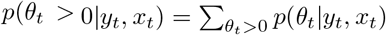
5. If *p*(*θ*_*t*_ *>* 0 | *y*_*t*_, *x*_*t*_) *>* 0.5, respond that the orientation is clockwise; if *p* < 0.5, respond counter-clockwise; if *p* = 0.5, respond randomly.
6. Compute the prior over contexts for the next time step; i.e., compute 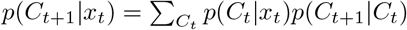

We used the following parameter values in the simulations: orientation = [−7.5, −5, −2.5, 0, 2.5, 5, 7.5], *σ*^2^ = 2 for high contrast stimuli and 5 for low contrast stimuli, *h* = 10, 20, and 50 %, *κ* = 0.4, 0.2, and 0.025 for Fig. 2e, and 0.45, 0.4, and 0.35 for Fig. S1. Every simulated experiment consisted of 57,600,000 trials.

### Signal Detection Theory model

We measured observers’ uncertainty about stimulus orientation and their estimation bias by fitting the relation between the task variables (orientation, contrast, and context) and probability of a “clockwise” choice with a Signal Detection Theory based process-model of decision-making ^35^. Under this model, each trial gives rise to an orientation estimate which is compared with a fixed criterion to obtain a decision (Fig. 3b). We assume that these estimates follow a Gaussian distribution, the mean of which is determined by the true stimulus orientation plus a context-specific constant (yielding two free parameters: one for the uniform context, and one for the non-uniform contexts). The spread of the Gaussian is determined by the contrast of the stimulus (resulting in two free parameters). Finally, we assumed that on some trials, observers “lapse” and simply guess without considering the task variables ^36^ (two free parameters, one per contrast). We chose this model based on a model comparison analysis in which we evaluated four versions of the model on the data collected during the test phase of the experiment (a 10,000-fold leave-one-out cross-validation analysis performed separately on high and low contrast choice data). Model versions differed in the number of free parameters used to describe the context-specific shift and spread of the orientation estimates (see Supplementary Table 2 and 3). Model parameters were optimized by maximizing the likelihood over the observed data, assuming responses arise from a Bernoulli process. We obtained confidence intervals on the uncertainty and bias estimates by performing a 1000-fold non-parametric bootstrap.

### Dynamic Bernoulli generalized linear model

We characterized the temporal evolution of our subjects’ decision-making strategy by fitting a descriptive model that specifies the trial-by-trial influence of a set of independently manipulated task variables (orientation, contrast, and context) as well as two history terms (previous response, previous orientation) and the nonlinearly transformed trials since context switch on the choice behavior. These predictors were linearly combined and then passed through a logit link function. Model parameters were optimized by maximizing the likelihood over the observed data, assuming responses arise from a Bernoulli process. We used a two-step “grid-search” procedure to find this maximum, whereby we first optimize all model parameters for a given value of *γ*, repeat this search-process for a manually specified range of *γ* values, and then select the solution with the best goodness-of-fit value. For each subject, the dynamic GLM better captured the data than the Signal Detection Theory model (Akaike Information Criterion comparison, see Supplementary Table 1 and Supplementary Fig. 1). To assess the necessity of the dynamic bias function, we conducted a cross-validation analysis in which trials that occurred between 1 and 4 trials after a context switch comprised the training set and trials that were the 5th or later after a context switch made up the test set. The full dynamic GLM out-performed a reduced “static” version that lacked the dynamic bias function in four of six veridical cue condition observers and three of six ambiguous cue condition observers (see Supplementary Table 4).

## Acknowledgements

This work was supported by US National Institutes of Health grants T32 EY021462 (J.A.C.), EY032999 (R.L.T.G.), the European Union’s Horizon 2020 research and innovation programme under the Marie Skłodowska-Curie Grant Agreement No. 754411 (W.F.M.), and the Howard Hughes Medical Institute (A.M.H.).

## Author contributions

J.A.C. and R.L.T.G. conceived and designed the study. W.F.M. and A.M.H. developed the Bayesian observer model. J.A.C. performed model simulations. Y.H.B. developed the psychophysics setup. J.A.C. collected human subject data. J.A.C. and R.L.T.G. analyzed data. J.A.C. and R.L.T.G. wrote the manuscript with input from W.F.M. and A.M.H.

## Competing Interests

The authors declare no competing interests.

## Supplementary information

**Figure S1.**
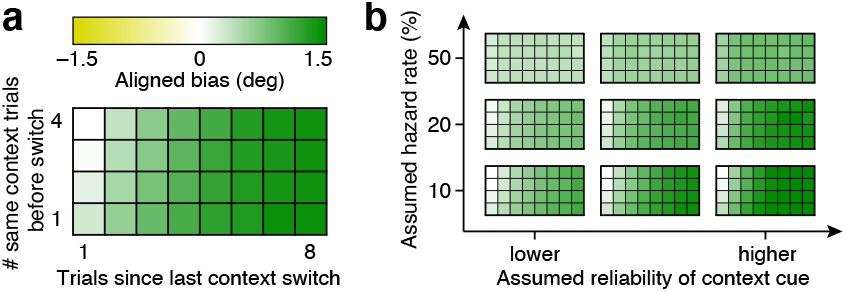
Determinants of the Bayesian ideal observer’s aligned bias. (a) Low-contrast aligned bias (color) as a function of trials since the most recent context switch (columns) and the number of same context trials *prior* to the most recent context-switch (rows) for an assumed hazard rate of 10% and a low level of context cue reliability. (b) The pattern of aligned bias across a range of assumed levels of hazard rate and context cue reliability.

### Comparison of Signal Detection Theory model and Dynamic GLM

We analyzed the choice data collected during the test phase of the experiment with two different models: a Signal Detection Theory based model (analyses in Fig. 3 and 4), and a dynamic GLM (analysis in Fig. 5). For each subject, the latter model provides a better description of the data. This is evident from the models’ AIC values, shown in Table S1 (the lower this value, the higher the quality of the model fit), and from a complementary analysis which compares each model’s normalized log-likelihood, split out for high and low contrast trials (Fig. S2).

**Figure S2.**
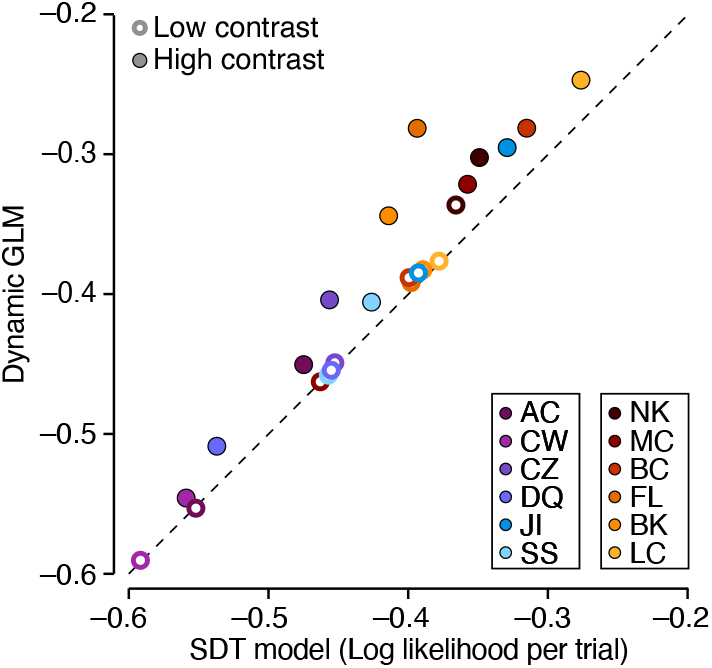
Comparison of goodness-of-fit of Dynamic GLM and Signal Detection Theory model. Low contrast data are shown as open symbols and high contrast data as filled symbols.

**Table S1.** AIC estimates for choice data collected during the test phase of the experiment under the Signal Detection Theory model and the Dynamic GLM.

| SDT vs GLM AIC values |  |  |
| --- | --- | --- |
| Subject | SDT | Dynamic GLM |
| AC | 5361.3 | 5238.7 |
| CW | 2804.1 | 2769.9 |
| CZ | 4932.9 | 4425.1 |
| DQ | 5179.0 | 4930.7 |
| JI | 3178.4 | 2995.8 |
| SS | 4059.8 | 3885.9 |
| MC | 4073.4 | 3894.6 |
| BC | 2035.9 | 1907.7 |
| BK | 3991.0 | 3608.8 |
| FL | 3479.9 | 2977.2 |
| LC | 3477.4 | 3308.6 |
| NK | 2414.1 | 2160.5 |

### Comparison of four Signal Detection Theory model variants

We considered four variants of the Signal Detection Theory model. These variants differed in the number of free parameters used to describe the context-specific shift and spread of the orientation estimates. Model 1 was the most economically parameterized variant by imposing a symmetric context-specific bias for the non-uniform contexts (resulting in two free “shift” parameters) and a single context-independent level of estimation uncertainty (resulting in one free “spread” parameter). Model 2 differed from this variant by allowing for a context-specific level of estimation uncertainty (yielding three free “spread” parameters), while Model 3 instead allowed for asymmetric context-specific bias (yielding three free “shift” parameters). Finally, Model 4 was the least restrictive variant by allowing asymmetric context-specific bias (three free “shift” parameters) and a context-specific level of estimation uncertainty (three free “spread” parameters). To evaluate each model’s performance, we trained the models on all trials except for three pseudo-randomly chosen hold-out orientations (one per context). We then computed the quality of the models’ prediction for this hold-out set. We repeated this procedure 10,000 times. As can be seen in the bottom row of Table S2 and S3, Model 1 tended to perform best.

**Table S2.** Average log likelihood for hold-out data predicted under four different variants of the Signal Detection Theory model (only high contrast trials included).

| High Contrast Data |  |  |  |  |
| --- | --- | --- | --- | --- |
| Subject | Model 1 | Model 2 | Model 3 | Model 4 |
| AC | -8.824 | -9.016 | -8.752 | -8.693 |
| CW | -7.821 | -8.353 | -7.973 | -8.515 |
| CZ | -10.335 | -10.995 | -10.613 | -11.355 |
| DQ | -10.106 | -10.622 | -10.234 | -10.426 |
| JI | -5.378 | -5.738 | -5.413 | -6.032 |
| SS | -9.987 | -10.331 | -10.190 | -10.457 |
| MC | -8.078 | -8.984 | -8.219 | -10.438 |
| BC | -6.506 | -6.146 | -7.197 | -6.658 |
| BK | -8.786 | -10.781 | -8.335 | -8.574 |
| FL | -6.626 | -13.649 | -6.290 | -10.015 |
| LC | -5.532 | -6.118 | -5.473 | -6.018 |
| NK | -5.992 | -5.659 | -6.237 | -5.872 |
| mean | -7.831 | -8.866 | -7.910 | -8.588 |

**Table S3.** Average log likelihood for hold-out data predicted under four different variants of the Signal Detection Theory model (only low contrast trials included).

| Low Contrast Data |  |  |  |  |
| --- | --- | --- | --- | --- |
| Subject | Model 1 | Model 2 | Model 3 | Model 4 |
| AC | -9.800 | -10.508 | -9.743 | -10.566 |
| CW | -7.669 | -7.974 | -7.825 | -8.205 |
| CZ | -6.975 | -7.907 | -6.775 | -6.969 |
| DQ | -5.955 | -6.263 | -6.195 | -6.543 |
| JI | -7.421 | -7.471 | -7.594 | -7.411 |
| SS | -7.848 | -10.646 | -8.301 | -10.856 |
| MC | -9.792 | -9.642 | -10.110 | -9.912 |
| BC | -7.434 | -8.315 | -7.582 | -8.401 |
| BK | -7.924 | -8.502 | -8.102 | -8.784 |
| FL | -7.560 | -7.798 | -7.275 | -7.334 |
| LC | -7.865 | -8.226 | -7.870 | -8.258 |
| NK | -7.171 | -7.288 | -7.066 | -7.396 |
| mean | -7.784 | -8.378 | -7.869 | -8.386 |

### Comparison of Dynamic and Static GLM

We analyzed the choice data collected during the test phase of the experiment with a GLM that included a dynamic bias function (analysis in Fig. 5). To assess the necessity of this model component, we conducted a cross-validation analysis in which we compared performance of this model with a variant that lacked this specific component (the “Static” GLM). These variants critically differ in the predictions they make for trials that occur after many trials within the same context. For this reason, we opted to use all trials that occurred between 1 and 4 trials after a context switch as training data and all trials that occurred 5 or more trials after a switch as hold-out test set. As can be seen in Table S4, the dynamic GLM outperformed the static version in four of six veridical cue condition subjects and three of six ambiguous cue condition subjects.

**Table S4.** Each cell of the table reports the total log-likelihood of the held out data for each subject. The rightmost column indicates the proportion of each subject’s total data the held-out fraction made up.

| GLM cross validation |  |  |  |
| --- | --- | --- | --- |
| Subject | Dynamic GLM | Static GLM | Percent data held out |
| AC | -364.822 | -365.256 | 14.2 |
| CW | -156.485 | -156.484 | 11.5 |
| CZ | -333.831 | -336.113 | 14.5 |
| DQ | -258.735 | -260.425 | 12.0 |
| JI | -201.178 | -201.000 | 13.0 |
| SS | -226.296 | -228.544 | 11.6 |
| MC | -254.774 | -255.147 | 12.6 |
| BC | -126.975 | -127.600 | 13.0 |
| BK | -244.683 | -245.215 | 14.6 |
| FL | -138.937 | -138.246 | 11.1 |
| LC | -259.729 | -259.425 | 14.5 |
| NK | -132.768 | -132.306 | 12.3 |

## Notes

### Competing Interest Statement

The authors have declared no competing interest.

